# Retinal microglial cells increase expression and release of IL-1β when exposed to ATP

**DOI:** 10.1101/2024.06.25.600617

**Authors:** Keith E. Campagno, Wennan Lu, Puttipong Sripinun, Farraj Albalawi, Aurora Cenaj, Claire H. Mitchell

## Abstract

Cytokine IL-1β is an early component of inflammatory cascades, with both priming and activation steps required before IL-1β release. Here, the P2X7 receptor (P2X7R) for ATP was shown to both prime and release IL-1β from retinal microglial cells. Isolated retinal microglial cells increased expression of *Il1b* when stimulated with endogenous receptor agonist extracellular ATP; ATP also rapidly downregulated expression of microglial markers *Tmem119* and *Cd206.* Changes to all three genes were reduced by specific P2X7R antagonist A839977, implicating the P2X7R. Microglial cells expressed the P2X7R on ramifications and responded to receptor agonist BzATP with robust and rapid rises in intracellular Ca^2+^. BzATP increased expression of IL-1β protein colocalizing with CX3CR1-GFP in retinal wholemounts consistent with microglial cells. ATP also triggered release of IL-1β from isolated retinal microglia into the bath; release was inhibited by A839977 and induced by BzATP, supporting a role for the P2X7R in release as well as priming. The IL-1β release triggered by ATP was substantially greater from microglial cells compared to astrocytes from the optic nerve head region. *Il1b* expression was increased by a transient rise in intraocular pressure and *Il1b* levels remained elevated 10 days after a single IOP elevation. In summary, this study suggests the P2X7 receptor can both prime IL-1β levels in microglial cells and trigger its release. The P2Y12R was previously identified as a chemoattractant for retinal microglia, suggesting the recruitment of the cells towards the source of released extracellular ATP could position microglia for P2X7R receptor, enabling both priming and release of IL-1β.

## Introduction

Microglial cells are critical for maintaining neuronal health, continuously monitoring the environment to reduce levels of cellular debris and removing redundant synapses [1, 2]. However, microglia are also involved in inflammatory signaling following disease-associated stimulation or autoimmune events [3]. Traditionally, microglial were thought to transition into distinct types of phenotypic states. However, emerging evidence suggests microglia can simultaneously exhibit a blend of protective and detrimental functions simultaneously, existing along a non-binary continuum that reflects their changing circumstances [4]. Transcriptional analysis has revealed microglia to be a heterogeneous population of cells, with location, age and environment contributing to their unique profiles [5–7]. Understanding how the beneficial and detrimental actions of microglia are balanced requires unraveling the complex factors that regulate microglial modulation.

IL-1β is a key cytokine released by microglia in inflammatory signaling [8, 9]. The release of IL-1β is tightly controlled, consistent with its pivotal role of in inflammation, and requires both priming and activation steps before the mature form of IL-1β is released from cells [10]. The cleavage and maturation of IL-1β is often mediated by the NLRP3 inflammasome, a multi-protein complex [11]. The NLRP3 inflammasome can be activated by multiple stimuli including lysosomal leakage of cathepsin enzymes and stimulation of the ionotropic P2X7 receptor for extracellular ATP [12]. Elevations in extracellular ATP capable of activation the receptor are associated with several neuropathologies where IL-1β is implicated such as traumatic brain injury and retinal degenerations including glaucoma [13–18]. This suggests that excess ATP acting on the P2X7 receptor could be an important step in the observed neuroinflammation. While responses of microglia in the brain have been studied in detail, considerably less is known about retinal microglial cells.

The present study examines the effects of extracellular ATP and the P2X7 receptor on IL-1β in retinal microglial cells and compares the release from microglia with that from astrocytes. The results suggest that stimulation of the receptor both primes IL-1β in microglial cells and triggers its release, highlighting the pathway’s significant impact on neuroinflammation.

## Methods

### Animals

All procedures were performed in strict accordance with the National Research Council’s “Guide for the Care and Use of Laboratory Animals” and were approved by the University of Pennsylvania Institutional Animal Care and Use Committee (IACUC) under protocol #803584.

Mice (C57Bl6/J, Cx3CR1-GFP, Jackson Laboratories (Bar Harbor, ME,) and Long–Evans rats (Harlan Laboratories/Envigo, Frederick, MD) of both sexes were used as sex was not considered a biological variable. All animals were housed in temperature-controlled rooms on a 12:12 light:dark cycle with food and water provided *ad libitum*.

### Microglial isolation

Primary retinal and brain microglia were isolated from mouse pups of both sexes (P12-P20) using the shake-off method as described [19, 20]. In both cases, mixed cell cultures were grown in media consisting of High Glucose DMEM (HG-DMEM; Invitrogen) with 10% Fetal Bovine Serum (FBS; Sigma-Aldrich), 1% Penicillin/Streptomycin (Pen/Strep; Gibco), 1% GlutaMAX (Gibco), and 1x MEM with nonessential amino acids (Sigma-Aldrich). Upon reaching confluence, microglia were gently “shaken off”, collected, and plated in dishes coated with Poly-L-Lysine (PLL; Peptides International, 0.01%) and Collagen IV (Corning, 4 µg/ml) in HG-DMEM with 5% FBS; inclusion of 10% of astrocyte conditioned media from the founding dish enhanced microglial development. Media was usually changed to DMEM with 5% FBS, 1% Pen/Strep, 1% GlutaMAX, and 1x MEM nonessential amino acids 24 hours prior to experimentation. Microglial cells were isolated from neonatal rat retinas using step 1 of the 2-step immunopanning protocol described previously [21]. In brief, retinas of Long Evans rat pups PD 3-7 of both genders were dissected from each eye globe and dissociated for 30 min at 37°C in Hank’s balanced salt solution (HBSS; Gibco, Inc. Invitrogen Corp., Carlsbad, CA) containing 15 U/mL papain, 0.2 mg/mL DL-cysteine and 0.004% DNase I. Retinas were washed and triturated in HBSS with 1.5 mg/mL ovomucoid, 1.5 mg/mL bovine serum albumin (BSA) and 0.004% DNase I, incubated with rabbit anti-rat macrophage antibody (10 min, 1:75, Accurate Chemical, Westbury, NY), centrifuged at 1000 rpm for 10 min, and washed. Cells were resuspended in phosphate-buffered saline (PBS) containing 0.2 mg/mL BSA and 5 µg/mL insulin, and incubated for 15 min in a 100 mm Petri-dish coated with goat anti-rabbit IgG antibody (1:400, Jackson ImmunoResearch Inc, West Grove, PA). After shaking and washing to remove unattached cells, microglia were detached with 0.05% trypsin and cultured with growth medium (HG-DMEM containing 10% fetal bovine serum (FBS), 1% Pen/Strep, 1% GlutaMAX, and 1x MEM nonessential amino acids) on 6-well plates.

### Astrocyte cultures

Primary cultures were grown from Long-Evans rat optic nerve head astrocytes as described [22]. In brief, rat pups of either sex were sacrificed by P5, and the optic nerve head was digested for 1-2 hours with 0.25% trypsin. After washing, cells were grown in DMEM/F12, 10% FBS, 1% Penicillin/Streptomycin, and 50 ng/ml epidermal growth factor (EGF; E4127, Sigma). Cells were used up to passage 5. Cultures were found to contain >99% astrocytes, as defined by glial fibrillary acidic protein (GFAP) immunofluorescence staining (MABH360, Chemicon International Inc).

### Retinal wholemounts

CX3CR1^+/GFP^ mice were euthanized by CO_2_ and their enucleated eyes were placed into an isotonic solution (consisting of (in mM) 105 NaCl, 5 KCl, 6 4-(2-hydroxyethyl)-1-piperazineethanesulfonic (HEPES) acid, 4 Na 4-(2-hydroxyethyl)-1-piperazineethanesulfonic acid, 5 NaHCO_3_, 60 mannitol, 5 glucose, 0.5 MgCl_2_, and 1.3 CaCl_2_, with a pH of 7.4). Retinal whole mounts were dissected as previously described [19]. Following exposure of live tissue to 2’(3’)-*O*-(4-Benzoylbenzoyl)adenosine-5’-triphosphate tri(triethylammonium) salt (BzATP) or isotonic solution for 6 hours at 37° C, retinas were fixed with 4% paraformaldehyde for 15 minutes at 25°C.

### Quantitative PCR

Retinas or isolated microglia were homogenized using TRIzol (Invitrogen). RNA was purified with an RNeasy mini kit (Qiagen, Inc.) and converted to cDNA using a High Capacity cDNA Reverse Transcription Kit (Applied Biosystems); qPCR was performed with PowerUp SYBR Green (Applied Biosystems) on the 7300 or Quant Studio 3 Real-Time PCR systems (Applied Biosystems) using standard annealing and elongation protocols, with data analyzed using the delta-delta CT approach as described [23]. Values are expressed as delta-delta CT as the exponential conversion to relative mRNA levels increases variation given the substantial rise in *Il1b* levels, although some examples have been converted in the text to provide context. Primers are as follows: *Il1b*: Forward: F: GGAGATTTCAAAAGCTGATGTGGA Reverse: R: CCTCAGACGCTGGTTGTCTT; Tmem119: F: GTGTCTAACAGGCCCCAGAA R: AGCCACGTGGTATCAAGGAG Tnfa: F: AAATGGCCTCCCTCTCATCAG R: GTCACTCGAA-TTTTGAGAAGATGATC Cd206 F: TCTTTGCCTTTCCCAGTCTCC R: TGACACCCAGCGGA-ATTTC *Gapdh:* F: TCACCACCATGGAGAAGGC R: GCTAAGCAGTTGGTGGTGCA.

### Ca^2+^ imaging

Intracellular Ca^2+^ was determined from microglial cells using previously described approaches [24]. The Fura-2 was chosen as a ratiometric Ca^2+^-indicator is preferable when BzATP induces morphological changes in microglial cells and rapid retraction of processes ([19], Fig S1). In brief, microglia were plated on 25 mm coverslips coated with PLL (0.01%) and Collagen IV (4 µg/ml) and loaded with 10 µM Fura-2 AM (Thermo Fisher) with 0.02% Pluronic F-127 (Thermo Fisher) for 45 min at 37°C. Cells were washed, mounted in a perfusion chamber, and perfused with isotonic solution without Mg^2+^, to avoid block of the P2X7R by Mg^2+^ [25]. Ratiometric measurements were performed using a 40x objective on a Diaphot inverted microscope (Nikon) by alternating excitation between 340nm and 380nm wavelengths with a scanning photometer and quantifying emission ≥512 nm with a charge-coupled device camera (All Photon Technologies International). Data were expressed as the ratio of light excited at 340nm to 380nm, F_340/380_, due to the complexity of calibration.

### IL-1β release

Microglial cells were primed with lipopolysaccharide (LPS; Sigma-Aldrich), or sometimes with IL-1α as indicated in the text, prior to agonist exposure. IL-1β release was measured using the Mouse IL-1 beta/IL-1F2 Quantikine ELISA kit (R&D Systems) following the manufacturer’s instructions, with samples, standards, and controls allowed to bind overnight at 4°C. Optical density was measured at 450 nm with subtraction of 540 nm measurements using the SpectraMax ABS (Molecular Devices). Values were converted back into absolute amounts using the standard curve. Due to the high variability between experiments, data were frequently normalized. Rat microglial cells and astrocytes were primed with 500 ng/ml LPS and rat IL-1α (R&D Systems) followed by agonist exposure. IL-1β was measured using the Rat IL-1 beta/IL-1F2 Quantikine ELISA Kit (R&D Systems) following the manufacturer’s instructions.

### Immunocytochemistry

Microglial cells were fixed in 4% paraformaldehyde for 10 min at 37°C, washed in PBS with 1% Tween 20 (Bio-Rad), permeabilized with 0.1% Triton-X 100 for 15 min (Sigma-Aldrich) and blocked with 20% Superblock (Thermo Fisher) plus 10% goat or donkey serum. For P2X7R/Iba1 staining, coverslips were incubated in anti-P2X7R polyclonal antibody (#APR-008, Alomone Labs, 1:200) and anti-Iba1 polyclonal antibody (#AB48004, Abcam, 1:200) overnight followed with with donkey anti-goat Alexa555 conjugated antibody (#A21432, Invitrogen, 1:500) and donkey anti-rabbit Alexa488 conjugated antibody (#A21206, Invitrogen, 1:500). For Iba1/GFAP staining used to determine purity, coverslips were incubated in anti-Iba1 polyclonal antibody (#019-19741, Wako, 1:500) and anti-GFAP monoclonal antibody (#MAB312 Chemicon, 1:500) overnight, followed by goat anti-rabbit Alexa546 conjugated antibody (#A11035, Invitrogen, 1:500) and goat anti-mouse Alexa488 conjugated antibody (#A11001, Invitrogen, 1:500). After incubation in Hoechst (Cell Signaling, 1 µg/ml) for 10 min, coverslips were washed and mounted using SlowFade Gold (Thermo Fisher). Wholemounts from Cx3CR1-GFP mice were permeabilized and stained for IL-1β using appropriate protocols [19]. Images were obtained using a Ti-2 Spinning Disc confocal microscope or an Eclipse inverted microscope using NIS Elements v. 4.6 Imaging software (all Nikon). ImageJ was used to modify intensity, and merge pseudocolored images, with control and experimental images processed in parallel.

### Transient Elevation of intraocular pressure (IOP)

The IOP was elevated mice using a modified version of the Control Elevation of IOP (CEI) protocol as described previously [19, 26]. C57BL/6J mice aged 3-6 months were anesthetized and maintained with 1.5% isoflurane throughout the procedure after receiving 5 mg/kg meloxicam. Corneal anesthesia and mydriasis were achieved by administering proparacaine (0.5%) and tropicamide (1%) eye drops, respectively. One eye was cannulated with a 33-gauge needle attached to polyethylene tubing (PE 50; Becton 51 Dickinson) inserted into the anterior chamber, connected to a 20 ml syringe filled with sterile hank balance salt solution (HBBS). IOP was elevated to 58–60 mmHg by positioning the reservoir to the appropriate height; this pressure was selected to maintain retinal blood flow and avoid ischemia Ocular lubricant gel was applied at regular intervals to both eyes to prevent cornea desiccation (GenTeal; Alcon laboratory, Fort Worth, TX). The needle was removed after 4 h and IOP returned to baseline, with 0.5% erythromycin ointment applied to the cornea. The contralateral eye served as a normotensive control.As a Sham control, the needle was inserted but the pressure was not elevated. Mice were euthanized 1 or 10 days later.

### Data analysis

Statistical analysis was performed using GraphPad Prism software v. 10 (GraphPad Software, LLC). Data were tested for normality using the Shapiro-Wilk or Kolmogorov-Smirnov test. Comparisons were performed with a one-way analysis of variance (ANOVA) or non-parametric test as appropriate, with Tukey’s test for comparisons between all samples and Dunnett’s test for comparisons to a control and Dunn’s test for comparisons for non-parametric tests. Results returning p<0.05 were considered significant. Data is represented as mean ± standard deviation. Observers were blinded to the experimental conditions wherever possible.

## Results

### Characterization of isolated microglial cells

Identifying the cellular source of cytokine released into a multicellular tissue is complex. To determine whether microglial cells released IL-1β, initial experiments utilized microglia isolated from the mouse retina. Isolated mouse retinal microglial cells had a ramified morphology (Fig. 1A). Several steps were taken to support the relevance of the cells to the in vivo situation. The responses to inflammatory agents LPS and IL-4 were characteristic of microglial cells. Specifically, a 4-hour exposure to LPS increased the expression of *Tnfa* mRNA while exposure to IL-4 for decreased *Tnfa* levels (Fig. 1B). LPS decreased levels of Tmem119 and CD206, while IL-4 increased the expression of both markers. This agrees with the responses previously described for Nos2 and Arg1 in brain and retinal microglia [19], strengthening the characterization. Immunohistochemical examination indicated the cells expressed monocyte marker Iba1 but not astrocyte marker GFAP (Fig. 1C); quantification indicated a microglial purity of >98% as described [19].

**Figure 1.**
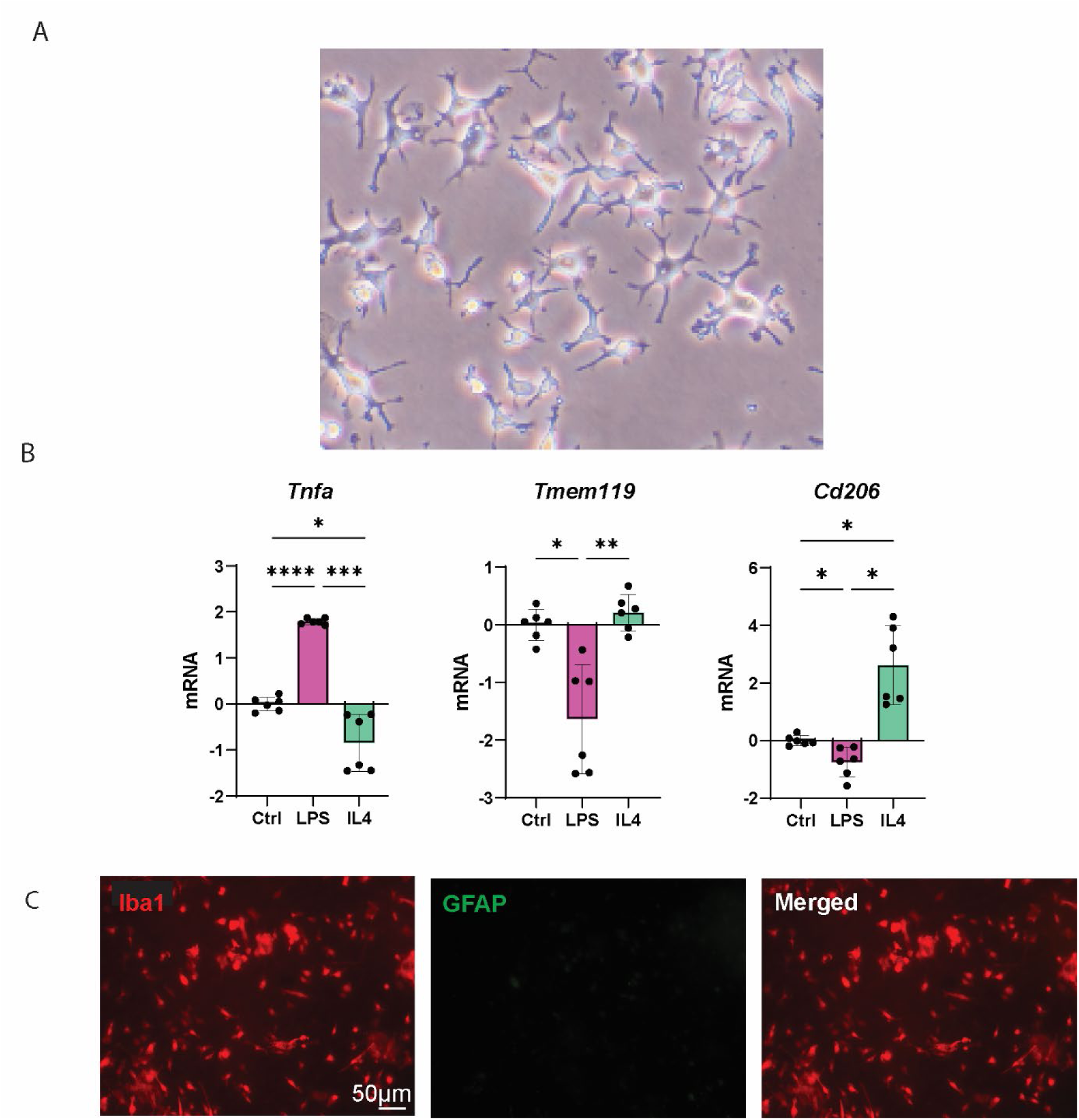
Characterization of isolated retinal microglial cells A. Representative image of isolated mouse retinal microglial cells. The ramified morphology was evident. B. qPCR results from isolated retinal microglial cells exposed for 4 hrs to DMSO (Ctrl), 10 ng/ml LPS (LPS), or 10 ng/ml IL-4 (IL4), showing changes in relative expression of mRNA for *Tnfa, Tmem119,* and *Cd206;* mRNA levels expressed as ΔΔCT values; dots represent the values from two replicates in triplicate, bars show the mean ± SD; *p<0.05, **p<0.01, ***p<0.001, ****p<0.0001. C. Immunohistochemical staining indicating the isolated microglial cells expressed microglial/monocyte marker Iba1 but not astrocyte marker GFAP.

### Microglial cells express the P2X7R and need priming to express Il1b

Immunohistochemistry with an antibody raised against an extracellular epitope confirmed the presence of the P2X7 receptor in cells with protrusions and expressing Iba1 (Fig. 2A). As the P2X7 receptor is an ionotropic cation channel with considerable permeability to Ca^2+^ [27], functional expression of the receptor was confirmed by measuring the ability of receptor agonist BzATP to elevate cytoplasmic calcium. BzATP induced a robust, reversible and repeatable elevation of intracellular Ca^2+^ (Fig. 2B). Together, these observations support the presence of the P2X7 receptor on these cells.

**Figure 2.**
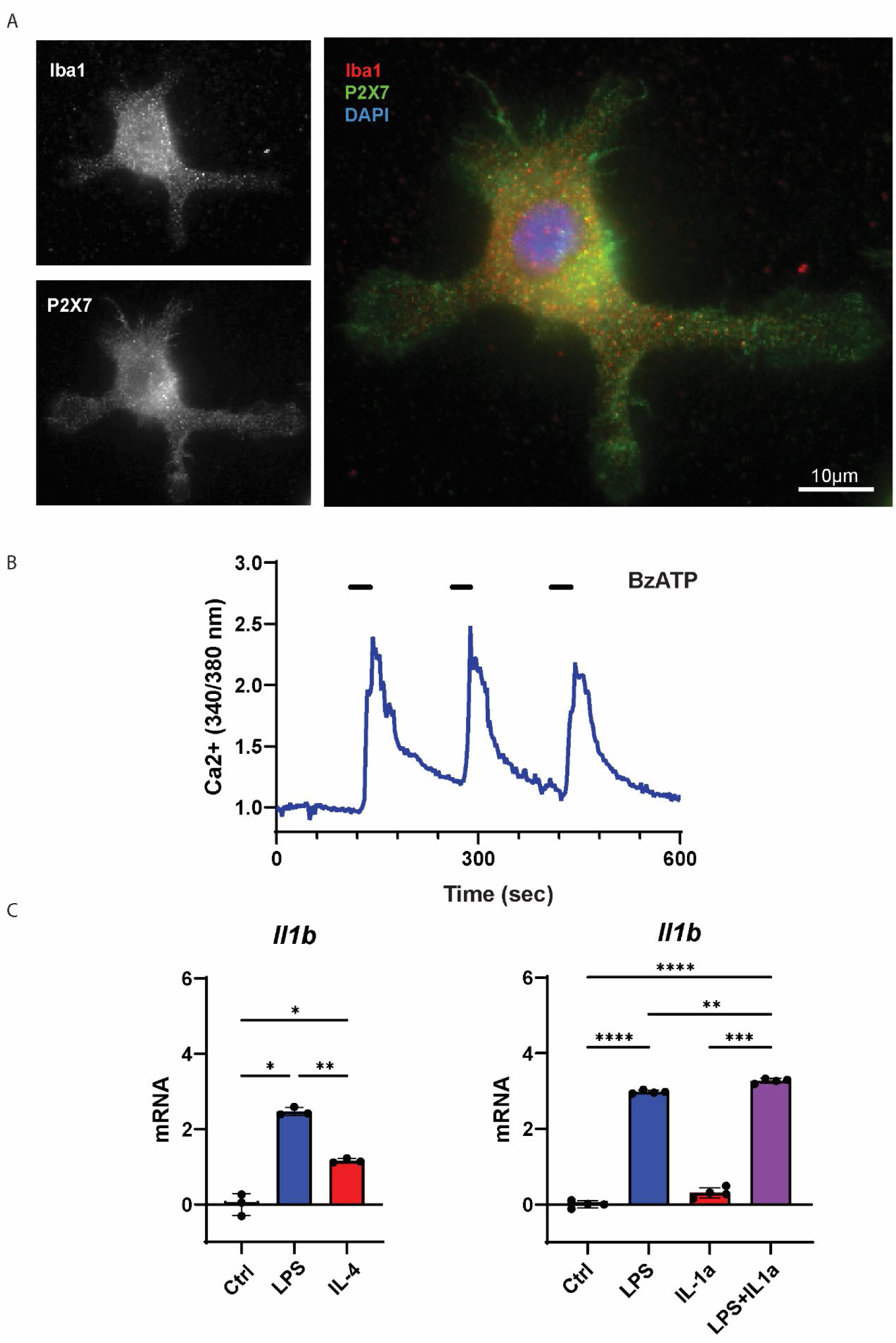
Microglial cells express the P2X7R and need priming to express Il1b A. Immunohistochemical staining indicating cells expressing Iba1 are positive for the P2X7 receptor. B. The microglial cells responded to P2X7R agonist BzATP (100 µM) with a rapid, reversible and repeatable rise in cytoplasmic Ca^2+^ consistent with the activation of the P2X7 receptor. Similar responses were observed in >20 cells from 4 preparations. C. qPCR results from isolated retinal microglial cells exposed for 4 hrs to DMSO (Ctrl), 10 ng/ml LPS (LPS), or 10 ng/ml IL-4 (IL4), showing changes in relative expression of *IL1b*. D. IL-1a (200ng/ml) had no significant effect on its own but increased expression of IL1b when given concurrently with LPS (500 mg/ml) for 4 hours to brain microglial cells. For C and D, mRNA levels expressed as deltadeltaCT values; bars show the mean ± SD; *p<0.05, **p<0.01, ***p<0.001, ****p<0.0001.

IL-1β is tightly regulated, and an initial priming step is usually required to increase its expression and that of other NLRP3 inflammasome components involved in its processing [28]. Although levels of *Il1b* was low under control conditions, they increased substantially after a 4-hour exposure to LPS (Fig. 2C); expression increased over 5-fold when converted from ΔΔTC values. Interestingly, *Il1b* expression was also increased by IL-4, although the rise was less than observed for LPS. The ability of both agents to increase IL1b levels supports the current view that polarization of retinal microglia is not entirely binary [4].

### Stimulation of P2X7 receptor triggers IL-1β release

The ability of ATP to trigger IL-1β release from mouse microglia was investigated. Previous reports indicate that extracellular ATP can trigger IL-1β release from brain microglial cells after they have been primed with LPS [20], so initial experiments tested the need for both the priming and activation stages using an ELISA for IL-1β. Levels of IL-1β released into the bath were relatively low after exposure of microglial cells to LPS + IL-1α for 3 hours. For the next hour, cells were exposed to either fresh priming solution, or priming solution + 3 mM ATP. While the additional hour in fresh priming solution led to a modest rise in secreted IL-1β, levels were substantially increased in the presence of ATP (Fig. 3A). This confirmed that both the priming and activation steps were needed for IL-1β release.

**Figure 3.**
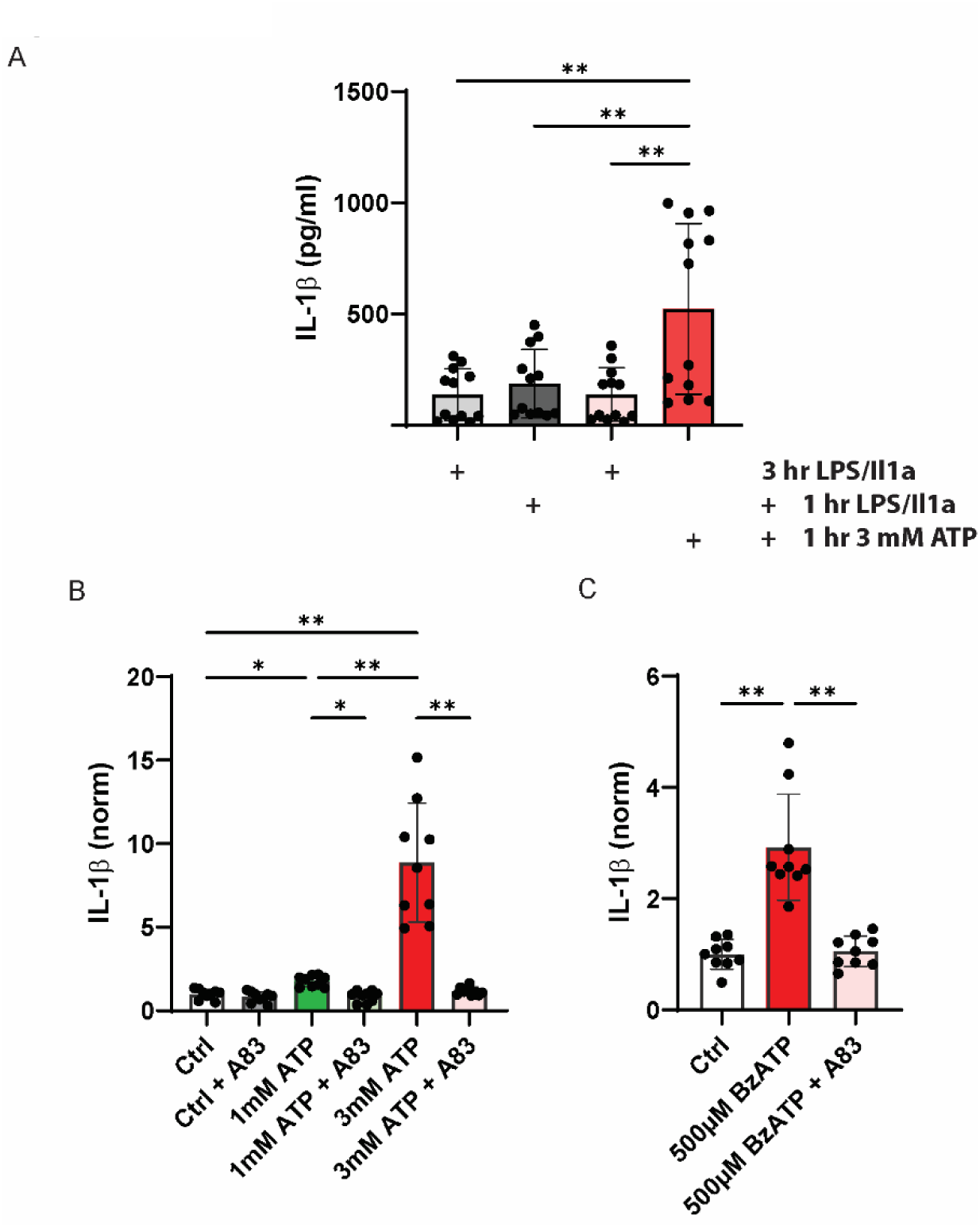
P2X7 receptor stimulation releases cytokine IL-1β from isolated mouse microglial cells A. ATP induces release of IL-1β from primed microglial cells. Retinal microglial cells were exposed to 1µg/ml LPS and 20ng/ml rrIL-1a for three hours, followed by an hour in the same priming solution or the solution plus 3 mM ATP. n=12; dots represent concentration from 4 independent experiments with 3 trials each. While the absolute levels of IL-1β sampled from the bath were low, this reflects the relatively large volume of extracellular space relative to cell volume. B. IL-1β release from microglial cells activated by 1 or 3 mM ATP was blocked by 1 µM P2X7 antagonist A839977. Microglia primed for 3 hours then ATP+priming solution added for 1 hour before collected as above. Data represent the mean ±SD from three wells each of 3 independent trials. As there was some variability in the absolute levels of IL-1β secreted from trial to trial, levels were normalized. C. P2X7R agonist BzATP (500 µM) induced IL-1β release from retinal microglial cells that was blocked by A839977. (3 independent experiments with 3 trials each). IL-1β levels normalized to the mean control values for each trial in B and C; *p<0.05, **p<0.01, ***p<0.001, ****p<0.0001.

The ability of the P2X7 receptor to trigger IL-1β release was tested next. A one-hour exposure to ATP in primed cells led to a dose-dependent rise in secreted IL-1β, with 1 mM ATP increasing levels by 77%. Although this concentration is sufficient to maximally stimulate other P2Y and P2X receptors, 3 mM ATP induced a much larger release, increasing levels by over 700% (Fig. 3B). The increased response with 3 mM ATP is consistent with the involvement of the P2X7 receptor [29]. This identification was supported by the ability of 1 µM A839977 to produce a complete block of the IL-1β released by 1 mM ATP and a 98% block of the release triggered by 3 mM ATP, as A839977 has a pIC_50_ = 6.8 at the mouse P2X7R [30].

Finally, the ability of P2X7R antagonist BzATP to trigger IL-1β release was evaluated. Exposure of primed cells to BzATP for one hour led to IL-1β secretion that was completely blocked by A839977. The dose response, agonist and antagonist responses strongly supported a role for the P2X7 receptor in the release.

### Priming of IL-1β by ATP in microglia

While the P2X7 receptor is most closely associated with the activation of the NLRP3 inflammasome, the receptor is also involved in the initial priming stage, leading to increased expression of several genes and proteins required for the assembly and processing of the IL-1β before release [31]. Our previous work demonstrated that ATP (and BzATP) increased expression of IL-1β on the mRNA and protein levels in astrocytes [31], so the presence of a similar priming event was examined in isolated mouse microglia. Exposure to 1 mM ATP alone for 4 hours increased the expression of *Il1b* message in microglia (Fig. 4A). This corresponds to a 20-fold rise in mRNA levels. ATP exposure also decreased expression of *Tmem119* and *Cd206* by approximately 4-fold. The effect of ATP on expression of all genes was reduced by 1µM A 839977, implicating the P2X7 receptor in the expression changes mediated by ATP.

**Figure 4.**
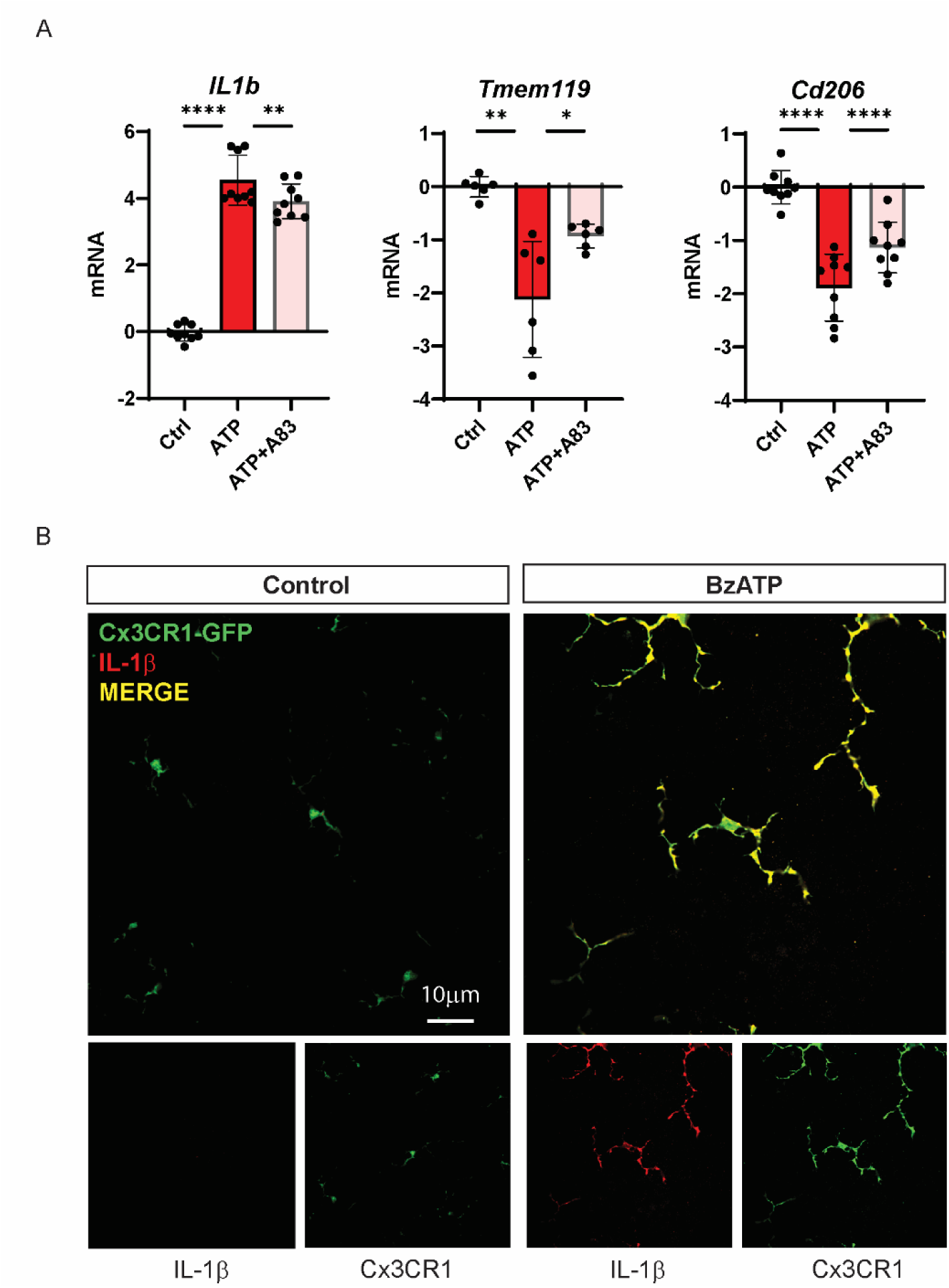
P2X7 receptor primes Il-1β in microglial cells A. Exposure of microglial cells to 1 mM ATP for 4 hours increased expression of *IL1b*, while the presence of P2X7 receptor antagonist A839977 (1µM) reduced the response. ATP decreased the expression of Tmem119 and Cd206, with A839977 reducing the response to ATP in each case. Data expressed as ΔΔCT. *p<0.05, **p<0.01, ***p<0.001, ****p<0.0001. B. Image of retinal whole mounts from Cx3CR1-GFP mice exposed to control media left) or 200 µM BzATP for 6 hours and immunostained for IL-1β (red).IL-1β was only detected in tissues exposed to BzATP, while the strength of the GFP signal (green) was increased by BzATP treatment. The overlap between IL-1β and GFP (yellow) was high. Representative of 11-14 regions from 4 mice..

### P2X7 receptor stimulation increases IL-1β on microglia of the Cx3CR1 mouse

While the ability of ATP to increase expression of Il1b mRNA in microglial cells, and of A839977 to block this, implicates the P2X7 receptor in priming, this needed to be confirmed at the protein level and in microglial cells integrated within the in vivo environment. Cx3CR1-GFP mice were used as the fluorescent labeling of monocytes enhanced microglial identification. Retinal whole mounts were bathed in agonist BzATP for 6 hours; trials indicated this exposure time enables diffusion of BzATP throughout the tissue. Under control conditions, moderate levels of GFP-positive cells were detected predominantly in the outer nuclear layer, inner limiting membrane and inner nuclear layer as expected. Little IL-staining of IL-1β was detected on Cx3CR1-GFP positive cells under control conditions, consistent with the low expression of *Il1b*. Exposure to BzATP increased levels of GFP-positive cells substantially, with increased expression surrounding the optic nerve as well as the throughout the retinal layers (Fig. 4B); the response reflected a response from withing the tissue as the use of explant prevented recruitment of Cx3CR1+ cells from elsewhere. BzATP increased the degree of IL-1β overlapping with Cx3CR1-GFP; expression of IL-1β overlapped with the Cx3CR1-GFP signal in the inner retina but was greater in the outer retina, and IL-1β expression was not restricted to microglial cells. This confirms that stimulation of the P2X7 receptor primes microglia by increasing expression of IL-1β at a mRNA and both protein level, and that this was present in intact tissue.

### ATP induces a larger release of IL-1β from microglial cells than astrocytes

Previous work has indicated that expression of *Il1b* in optic nerve head astrocytes was also increased following stimulation of the P2X7 receptor, but the ability of the astrocytes to release IL-1β was not clear [31]. Given the key role of astrocyte neurotoxicity in neurodegeneration [32], levels of IL-1β released from astrocytes with ATP exposure were determined and compared to the release from microglial cells. Experiments were performed rat cells as the isolation of rat optic nerve head astrocytes is more reliable, as this facilitates comparison with previous experiments, as it provides an additional species to increase experimental rigor [22]. Both astrocytes and microglia were primed for 3 hours to separate the effects of ATP on priming and release. A one-hour exposure to ATP increased the concentration of IL-1β bathing microglial cells over 7-fold but had no effect on the expression in astrocytes (Fig. 5).

**Figure 5.**
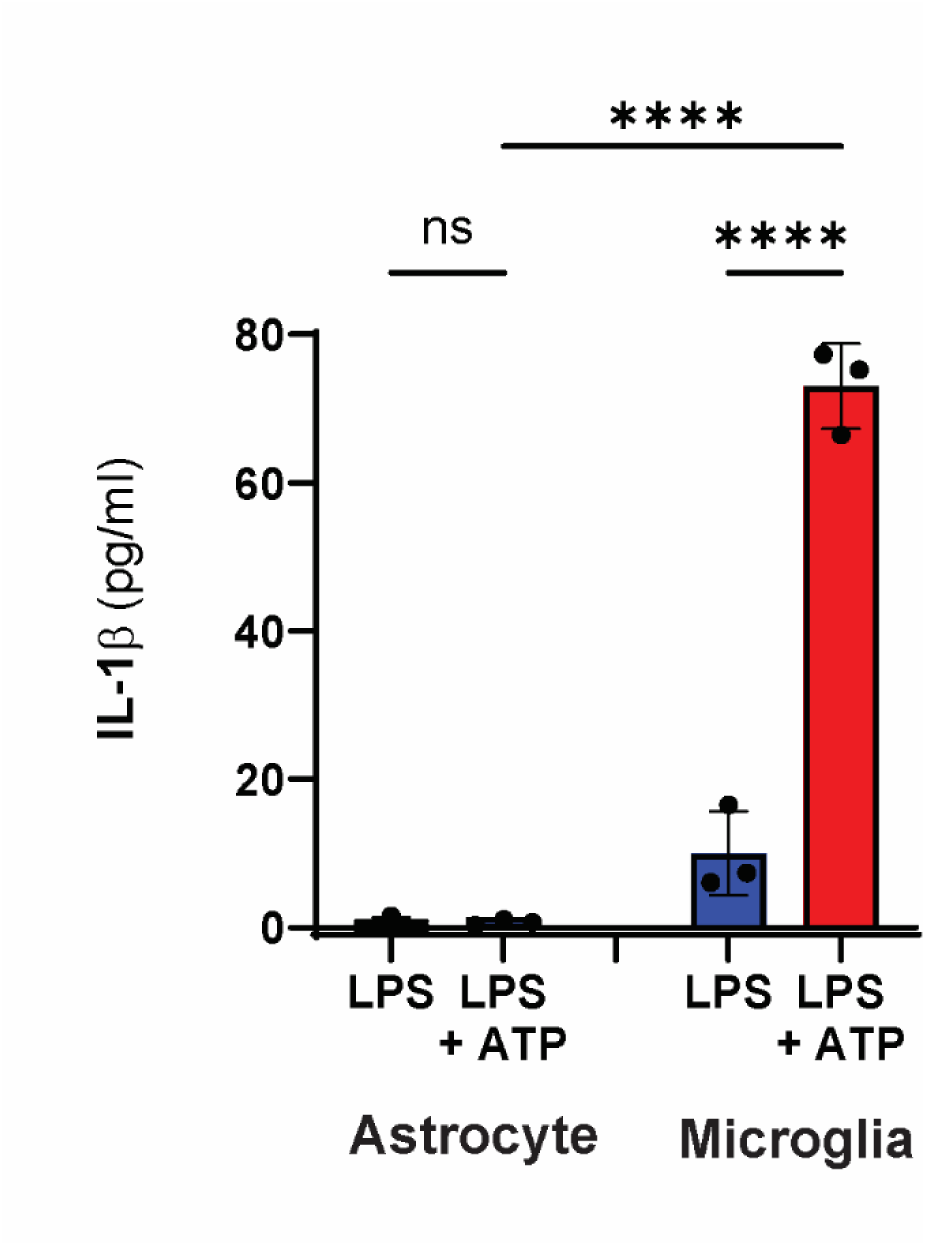
ATP induces a larger release of IL-1β from microglial cells than astrocytes Rat microglia primed with 500 ng/ml LPS for 3 hours followed by exposure to an additional 3 mM ATP released significantly more IL-1β than cultured rat astrocytes primed with LPS and 5 ng/ml IL-1α, followed by similar exposure to ATP (n=3) (ANOVA with Tukey’s multiple comparisons test.. ****p<0.0001.

### Sustained elevation of Il1b after single episode of IOP elevation

Given that cytokine signaling including IL-1β has been implicated in the pathogenesis of glaucoma, and that the elevation of IOP is a major risk factor for glaucoma, we asked whether elevation of IOP could increase expression of *Il1b* [33, 34]. A transient increase of IOP for 4 hours was sufficient to increase expression of Il1b (Fig. 6A-C); the values corresponded to >35-fold rise in *Il1b* expression. Interestingly, the Il1b expression remained elevated when examined 10 days after the IOP elevation. This suggests that a single episode of IOP elevation is sufficient to induce a sustained rise in *Il1b* levels.

**Figure 6.**
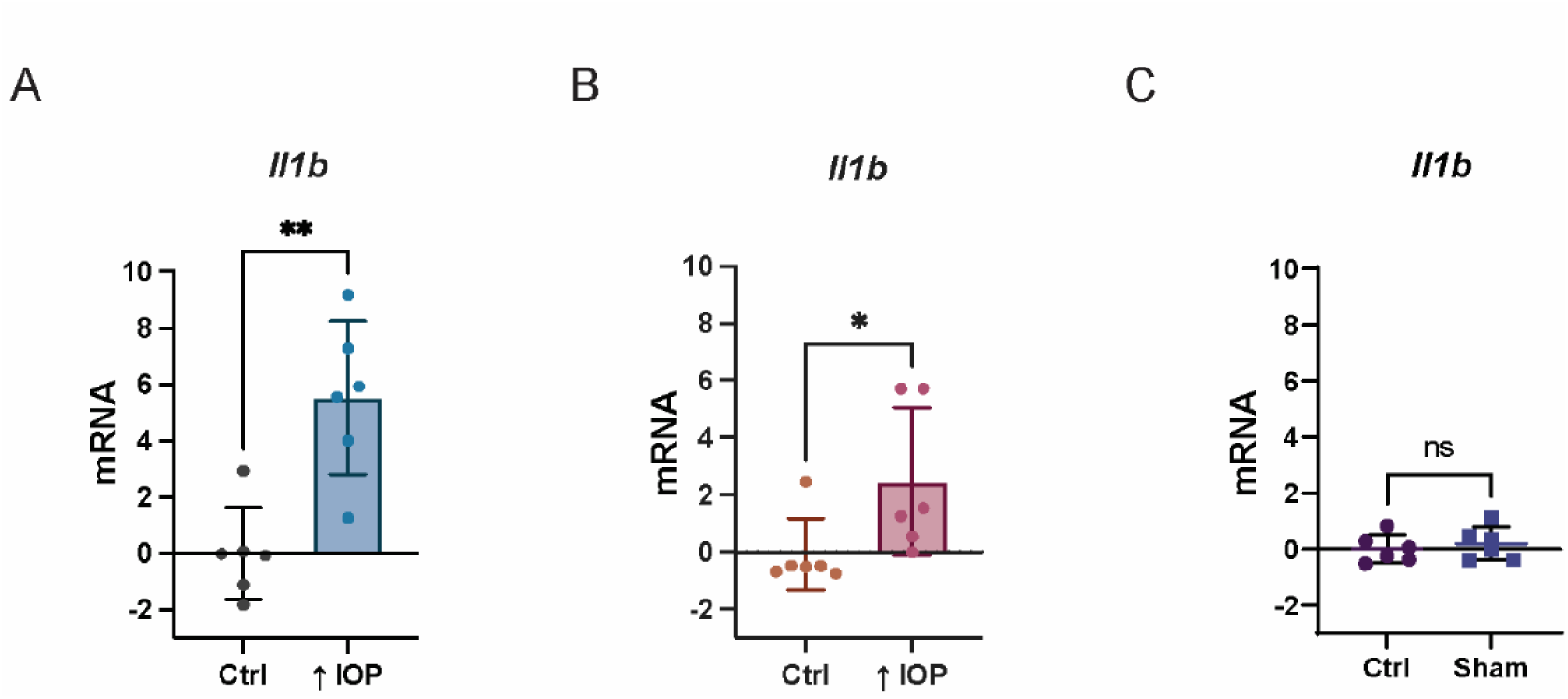
Sustained increase in Il1b levels with IOP rise A. Increased expression of retinal *Il1b* 1day after a transient 4-hour elevation of IOP (n=6, **p=0.0074). RNA expressed as ΔΔCT levels. B. *Il1b* remained elevated 10 days after a single IOP elevation (n=6, 0.031). C. Normotensive sham procedure did not alter *IL1b* expression (n=6, p=0.634).

## Discussion

This study examined the P2X7 receptor on retinal microglial cells in vivo and in vitro, focusing on its role in triggering the priming and release of IL-1β from these cells. The findings demonstrate that the P2X7 receptor is present on isolated mouse microglial cells both functionally and at the protein level. The release of IL-1β from isolated mouse microglial cells required an initial priming step, with LPS exposure increasing expression of IL-1β to enable a 1-hour exposure to extracellular ATP to trigger IL-1β release. The P2X7R antagonist A839977 reduced the IL-1β release induced by endogenous agonist ATP, while the more specific P2X7 agonist BzATP also evoked release, supporting receptor identification. A 4-hour exposure to extracellular ATP primed microglial cells, increasing the expression of *Il1b*. The P2X7 receptor also downregulated the expression of *Tmem119* and *Cd206*.Increased expression of IL-1β protein was also observed on microglial cells in Cx3CR1-GFP whole mounts after a 6-hour stimulation by BzATP, supporting the priming response in microglial cells withing the intact retina. The IL-1β release triggered by ATP was significantly greater from microglial cells compared to astrocytes from the optic nerve head region.

### IL-1β and the NLRP3 inflammasome

Previous studies have shown that transient non-ischemic elevation of IOP upregulates inflammasome components such as IL-1β, NLRP3, ASC, and CASP1 in mouse and rat retinas [31]. These findings were supported by the observation that BzATP injection mimicked the rise in *Il1b* and that this response was diminished in P2X7^-/-^ mice and by intravitreal injection of the P2X7 antagonist Brilliant Blue G. The current study supports these findings, suggesting that microglial cells significantly contribute to the observed changes in retinal tissue.

### Implications for microglial and astrocytic roles

Isolated optic nerve head astrocytes were previously shown to increase *Il1b* expression when subjected to swelling or stretching [31]. This effect was mitigated by P2X7R antagonists, removal of extracellular ATP, or blocking the transcription factor NFkB. Elevation of IOP also increased IL-1β immunostaining, with some overlap observed in GFAP-positive regions. Given these responses, the lower levels of IL-1β released from astrocytes as compared to microglial cells were somewhat unexpected. However, this discrepancy highlights the distinct roles of these cell types in the priming and activation steps of IL-1β release. The separation of priming and activation steps of IL-1β release between microglial cells and astrocytes highlights the complex regulation of inflammation in the retina, where microglial cells play a more prominent role in IL-1β secretion. The findings underscore the importance of microglial cells in the inflammatory response within the retina, particularly in the context of P2X7 receptor activation and IL-1β release, and agree with the association between macrophages and microglial cells and IL-1β release [31].

### Limitations of the current study

The use of isolated microglial cells brings some limitations, as microglial cells alter their characteristics when removed from their native environment [35]. However, these cells were characterized to show their relevance, including the ability of LPS to upregulate *Tnfa* and downregulate *Tmem119* and *Cd206*, and the ability of IL-4 to produce opposite changes. Additionally, the lack of GFAP expression was confirmed, supporting the purity of the microglial cell cultures. The predominance of IL-1β in outer retinal microglial cells as compared to inner retinal microglia and its presence in other retinal cells was also unexpected. While the data still support the idea that microglial cells are key contributors to IL-1β release in the retina, the exact mechanisms and the role of other retinal cells require further investigation.

The response observed with purinergic P2X7R overexpression in driving microglia activation and proliferation seems to be considerably greater than the rise in IL-1β observed in rat retinal microglial cells. Additionally, the release of IL-1β triggered by P2X7R overexpression alone in the hippocampal cultures appears to be more pronounced than what was observed in the retinal microglial cells, especially considering that the microglial cells in the retinal study were not primed with LPS or any other activating agent [36].

### Physiological relevance

The IL-1β released by retinal microglia is likely to have several detrimental effects on retinal health. For instance, studies have shown that IL-1β from microglia can increase chemokine levels in retinal pigment epithelial (RPE) cells and Müller cells, contributing to retinal degeneration [36]. Activation of the inflammasome in macrophages has also been associated with photoreceptor cell death [37], and IL-1β released from subretinal monocytes has been linked to cone cell death [38]. The current study identifies extracellular ATP in the retina as a possible signal to initiate these responses. Retinal ATP release can also occur as a cotransmitter from excitatory and inhibitory synaptic vesicles from horizontal, amacrine, and retinal ganglion cells, although the levels of ATP released in synapses are usually tightly controlled both spatially and temporally [39, 40]. ATP is released from astrocytes and RPE cells following stimulation of Toll-like receptors 3 and 4 [22] and from Muller cells stimulated with glutamate [41]. Increased release of ATP was detected in retinas from a murine model of Alzheimer’s disease as compared to controls [42]. The sustained increase in ATP release from mouse, rat and non-human primate models of chronic IOP elevation [43]. This increase in ATP may contribute to the observed rise in IL-1β levels, implicating a role in the pathophysiology of glaucoma [44], although further research is needed to directly confirm the role of sustained ATP release in mediating IL-1β increases in glaucoma. Regardless, the elevation of Il1b expression 1 and 10 days after a single 4 hour episode of IOP elevation suggests that fluctuations in IOP known to occur may impact the neuroinflammatory response [45].

Ultimately, the context in which IL-1β is released from microglia depends on the magnitude of ATP release and the proximity of microglial cells to the ATP source. The P2Y12 receptor plays a crucial role in recruiting microglia, bringing them closer to the ATP release source and exposing them to higher ATP concentrations. Most P2X and P2Y receptors respond to ATP at concentrations ranging from 30 to 100 µM [46, 47]. However, the potency of ATP acting on the mouse P2X7 receptor (P2X7R) is indicated by a pEC_50_ value of 2.6 [29], Although only two ATP concentrations were tested in the study, the observed response suggests the involvement of the P2X7R in IL-1β release from microglia.

### Conclusions

Overall, the results from the present study reinforce the significance of the P2X7 receptor in mediating IL-1β release from retinal microglia. Further research is needed to confirm the presence of IL-1β in other retinal cells and to elucidate the detailed mechanisms underlying the differential roles of microglial cells and astrocytes in retinal inflammation. Understanding these pathways could provide new therapeutic targets for managing retinal inflammatory conditions.

